# Impact of perinatal maternal docosahexaenoic acid-containing phospholipid synthesis on offspring growth and neurological symptoms

**DOI:** 10.1101/2024.01.06.574487

**Authors:** Ayumi Kanatani, Daisuke Hishikawa, Katsuyuki Nagata, Fumie Hamano, Kenta Nakano, Tadashi Okamura, Takao Shimizu, Hideo Shindou, Takeshi Nagamatsu, Keisuke Yanagida

## Abstract

Mothers provide essential nutrients, including docosahexaenoic acid (DHA), an omega-3 fatty acid, during the perinatal period. DHA deficiency in perinatal mothers is linked to developmental abnormalities, especially in the central nervous system of the offspring; however, its specific impact on distinct events in fetal and neonatal brain development and prospective brain functions remains incompletely understood. We demonstrated using mice lacking *Agpat3*, a gene encoding the enzyme that synthesizes DHA-containing phospholipids (DHA-PLs), that maternal DHA-PL synthesis significantly contributes to the maternal– offspring DHA supply during the fetal period but not in infancy. Selective modulation of DHA-PL levels during fetal and postnatal periods in *Agpat3*-knockout mice showed that fetal stage-specific insufficiency in maternal DHA-PL supply potentially influences the neuropsychiatric phenotype in adult mice without affecting postnatal tissue DHA-PL levels, weight gain, and brain expansion. Collectively, enhancing maternal DHA-PL synthesis during pregnancy may help prevent prospective neuropsychiatric abnormalities in the offspring.

## Introduction

Omega-3 fatty acids are pivotal nutrients for brain development and function ^1, 2^. Docosahexaenoic acid (DHA) is the most abundant omega-3 fatty acid within the brain, primarily existing as a component of membrane phospholipids (PLs) ^3, 4^. DHA-containing phospholipids (DHA-PLs) play diverse roles in brain development and function, serving as cellular membrane components and a reservoir of non-esterified DHA ^5^. The enrichment of DHA-PLs in neuronal membranes is hypothesized to support functions such as neurotransmission and neurite/axonal outgrowth by regulating cellular membrane biophysical properties ^6, 7^. Once released from membrane PLs, DHA and its derivatives govern neuronal cell differentiation, functions, and inflammation by acting as ligands for plasma membrane and nuclear receptors, thereby regulating intracellular signaling and transcription ^5, 8^.

Omega-3 fatty acid intake is particularly crucial during the perinatal period marked by rapid brain development in fetal and neonatal stages ^9^. Animal models and human intervention studies consistently demonstrate that omega-3 fatty acid deficiency during this critical period adversely affects brain development, leading to neuropsychiatric abnormalities ^10, 11, 12, 13^. Notably, processes such as neuronal migration, neurogenesis, and microglia invasion predominantly occur in fetal stages, whereas experience-dependent synaptic pruning becomes more prominent after birth ^14^. The specific roles of DHA in fetal and neonatal brain development must be elucidated to comprehend the mechanisms underlying neuropsychiatric abnormalities triggered by perinatal omega-3 fatty acid deficiency.

Fetuses and infants depend on maternal DHA supply since mammals lack the ability to synthesize omega-3 fatty acids *de novo*. Infants are presumed to receive DHA from breast milk fat, primarily composed of triglycerides (TGs) ^15^. Conversely, fetuses obtain DHA from various types of DHA-containing lipids in maternal blood via the placenta. The elevation of DHA-containing lipids in maternal serum during gestation ^16, 17, 18^ and the preferential transfer of DHA over other fatty acids ^18, 19^ highlight the importance of maternal DHA supply for fetal development. However, our understanding of the mechanisms governing maternal–fetal DHA transfer remains incomplete, and there is a notable lack of experimental evidence regarding the respective contributions of DHA-PLs and/or DHA-TGs in maternal blood.

The incorporation of DHA into phosphatidic acid (PA, a common precursor for PL synthesis), by 1-acyl-*sn*-glycerol-3-phosphate acyltransferase 3 (AGPAT3, also termed lysoPA acyltransferase 3 or lysoPL acyltransferase 3) is pivotal for maintaining cellular DHA-PL levels ^20, 21, 22^. Loss of *Agpat3* in mice results in a substantial decrease in tissue and plasma DHA-PL levels, without apparent impact on DHA-TG levels ^23^. This study utilized the *Agpat3*-knockout (KO) mouse model to investigate the mechanisms of maternal–fetal DHA transport during the perinatal period and its implications for brain development and functions. Furthermore, we explored the consequences of DHA deficiency during the fetal and infant periods using this model. Collectively, we believe that our findings would help elucidate the impact of fetal stage-specific DHA-PL insufficiency on neuropsychiatry in adult mice and may have broader implications for interventions targeting omega-3 fatty acid deficiencies during crucial developmental periods.

## Results

### Docosahexaenoic acid-containing lipid levels in maternal blood and fetal tissues increase during gestation

The molecular basis of maternal DHA supply to fetal tissues during gestation was elucidated by assessing the time-dependent lipidomic changes in pregnant wild-type (WT) C57BL/6J mouse serum using liquid chromatography-tandem mass spectrometry (LC-MS/MS). We monitored the levels of the major DHA-containing lipid species in blood [TGs, cholesterol ester (CE), phosphatidylcholine (PC), and lysoPC (LPC)] on gestational days 11 (GD11), GD14, and GD18 (Fig. 1a and Supplementary Fig. 1a–e). The levels of DHA-TGs and DHA-containing LPC remained unaltered during this period (Fig. 1e, f, and Supplementary Fig. 1c–f). Conversely, the relative abundances of PC 38:6 and PC 40:6 (typically the most abundant DHA-PC species), and DHA-CE increased with the progression of pregnancy (Fig. 1b–d and Supplementary Fig. 1a, b). This suggests that maternal synthesis of DHA-PLs or -CE during pregnancy likely contributes to DHA supply to fetal tissues in the late stage of pregnancy.

**Fig. 1.**
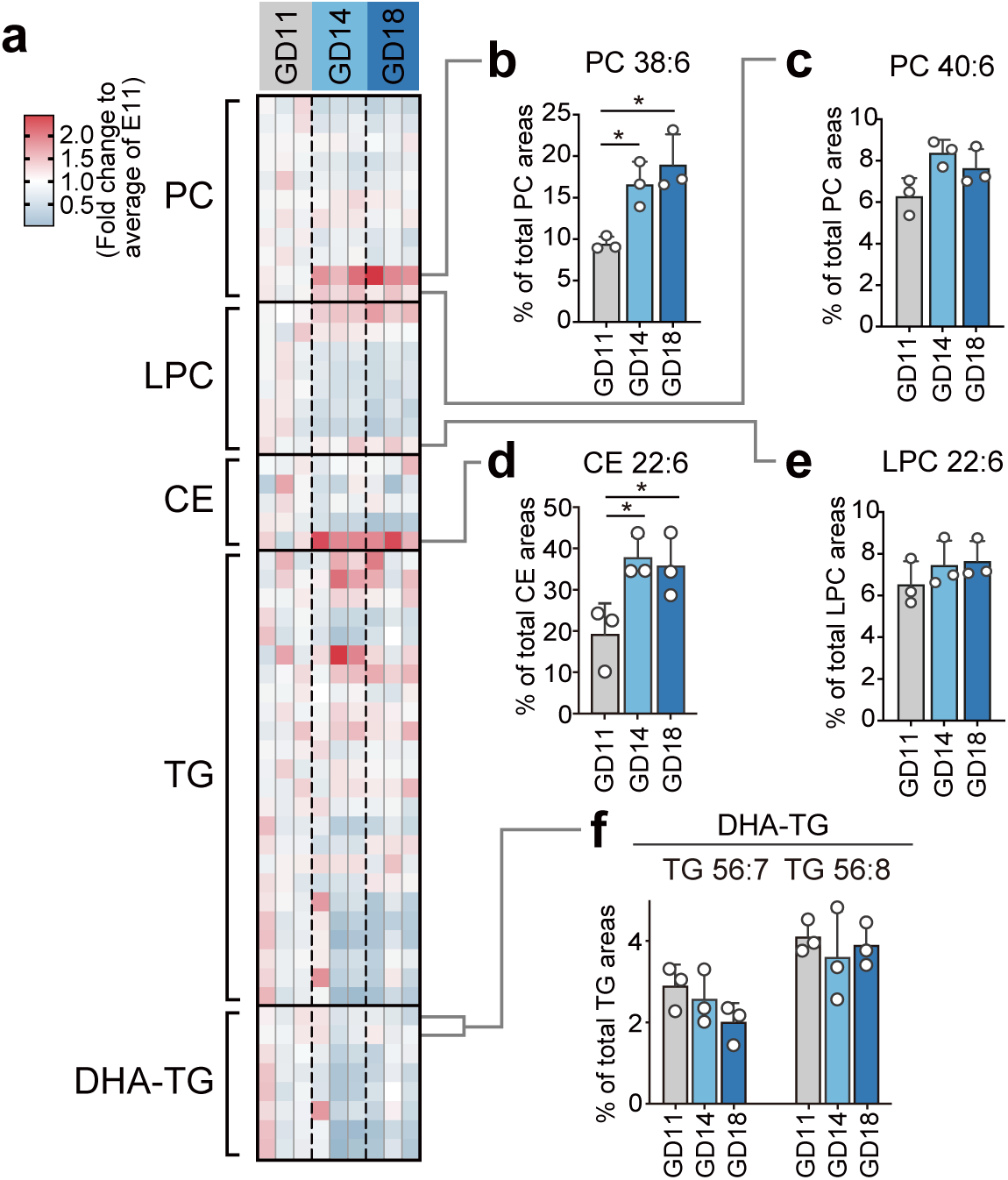
Changes in serum lipid profiles during pregnancy. (a) Comparison of phosphatidylcholine (PC), lysoPC (LPC), cholesterol ester (CE), and triglyceride (TG) levels at different gestational days (GD11, GD14, and GD18) based on their relative abundance within each lipid class. Heatmap depicting fold changes relative to mean levels on GD 11 for each lipid species (n = 3). (**b**–**f**) Relative abundances of docosahexaenoic acid (DHA)-containing PC (DHA-PC) (**b** and **c**), DHA-CE (**d**), DHA-LPC (**e**), and DHA-TG (**f**). Data are presented as means + SD. (n = 3 for each group). Statistical significance was determined using Bonferroni’s multiple comparisons (\**P* < 0.05).

In parallel with the increases in DHA-PCs and -CE in maternal blood, fetal tissue levels of DHA-PLs (such as PC and phosphatidylethanolamine (PE)) increased during gestation (Supplementary Fig. 2). In addition to the direct DHA supply from the mother, conversion of DHA from omega-3 fatty acid precursors, such as α-linolenic acid (FA 18:3, n-3) and eicosapentaenoic acid (EPA; FA 20:5, n-3), in fetal tissues may contribute to the increase in fetal DHA-PL levels during development. This conversion is catalyzed by fatty acid elongases (ELOVL2 and ELOVL5) and fatty acid desaturases (FADS1 and FADS2) and is thought to exclusively occur in the liver ^24^. Quantitative reverse transcription (RT-q) PCR analysis indicated that the mRNA levels of these enzymes were substantially low in fetal tissues compared with those in the neonatal mouse liver (Supplementary Fig. 3). Thus, DHA levels in fetal tissues likely depend on maternal DHA supply rather than *de novo* production from their precursors.

### Manipulation of maternal DHA-PL levels alters fetal DHA-PL levels

We recently reported that hepatocyte-specific *Agpat3* deletion led to decreases in the levels of DHA-PLs, -lysoPL, and -CE without affecting DHA-TG levels in mouse plasma ^23^. Building upon these findings, we investigated the contribution of maternal DHA-PL and DHA-TG synthesis to maternal–fetal DHA transfer using global *Agpat3*-KO mice.

There were marked decreases in the proportions of DHA-PCs, -LPC, and -CE in the serum of pregnant *Agpat3*-KO mice, whereas no decrease was observed in the proportion of DHA-TGs and the total TG amount according to LC-MS/MS analysis (Fig. 2). The DHA-CE levels in the plasma of hepatocyte-specific *Agpat3*-KO mice were markedly reduced, although the levels in the liver were unaltered ^23^. This is likely because most circulatory CEs are formed by fatty acid transfer from PC to free cholesterol in the blood through the action of lecithin cholesterol acyltransferase ^25^. Therefore, the reduction in plasma DHA-CE levels in *Agpat3*-KO mice appeared to be secondarily derived from DHA-PC deficiency in the blood. Collectively, preventing DHA-PL synthesis in pregnant mice decreased circulatory DHA-containing lipid levels but not DHA-TG levels.

**Fig. 2.**
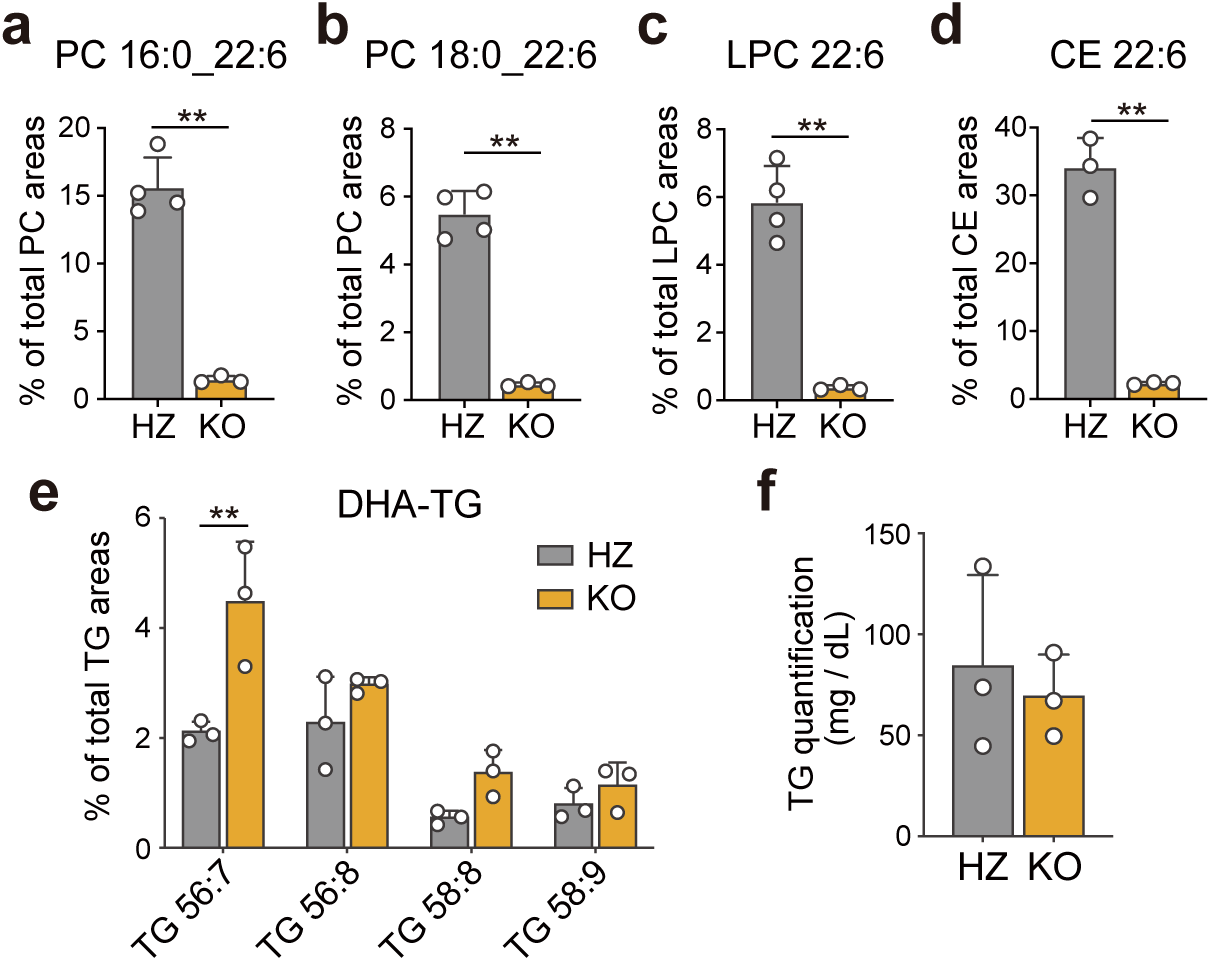
Levels of docosahexaenoic acid (DHA)-containing lipids and triglycerides in *Agpat3*-HZ and -KO mouse sera. (**a**–**e**) Relative abundances of DHA-containing phosphatidylcholines [DHA-PC; PC 16:0_22:6 (**a**) and PC 18:0_22:6 (**b**)], -lysoPC (DHA-LPC) (**c**), -cholesterol ester (DHA-CE) (**d**), and - triglycerides (DHA-TGs) (**e**) in *Agpat3*-HZ and -KO mouse sera (**a**–**c**, n = 4 for HZ and n = 3 for KO; **d** and **e**, n = 3 for each genotype). (**f**) Serum TG level in *Agpat3*-HZ (n = 3) and -KO (n = 3) mice. Data are represented as means + SD (n = 3 for each group). Statistical significance was determined using an unpaired *t*-test (\*\**P* < 0.01) for **a**–**d** and **f**, and using Bonferroni’s multiple comparisons (\*\**P* < 0.01) for **e**.

We hypothesized that the contribution of maternal DHA-PL synthesis to fetal DHA transfer could be validated by examining DHA levels in the offspring of *Agpat3*-KO based on lipidomic profiles of pregnant *Agpat3*-KO mouse plasma. Therefore, we prepared fetuses with different maternal DHA-PL levels by crossing *Agpat3* heterozygous (-HZ) or -KO females with *Agpat3*-HZ male mice (Fig. 3a). A comparison of *Agpat3*-HZ and -KO fetuses born from *Agpat3*-HZ dams showed that the DHA-PL levels in the *Agpat3*-KO placenta and fetal liver, brain, and plasma decreased by approximately 80%, 70%, 40%, and 70%, respectively, compared with those in the *Agpat3*-HZ fetuses (Fig. 3b–e, light gray versus dark gray). Similar trends were observed in *Agpat3*-HZ and -KO fetuses born from *Agpat3*-KO dams (Fig. 3b–e, green versus magenta). These limited declines were in sharp contrast to the drastic reductions (to <10% of WT) in DHA-PL levels in various tissues of adult and neonatal mice ^23, 26, 27^. Therefore, fetal tissues likely maintain considerable DHA-PL levels even in the absence of their AGPAT3 enzymes unlike postnatal tissues. Notably, the maternal *Agpat3* genotype significantly affected the DHA-PL levels in the fetuses, irrespective of the fetal genotypes (Fig. 3b–e, light gray versus green and dark gray versus magenta). These data suggested that maternally synthesized DHA-PLs are the key determinant of fetal DHA-PL levels.

**Fig. 3.**
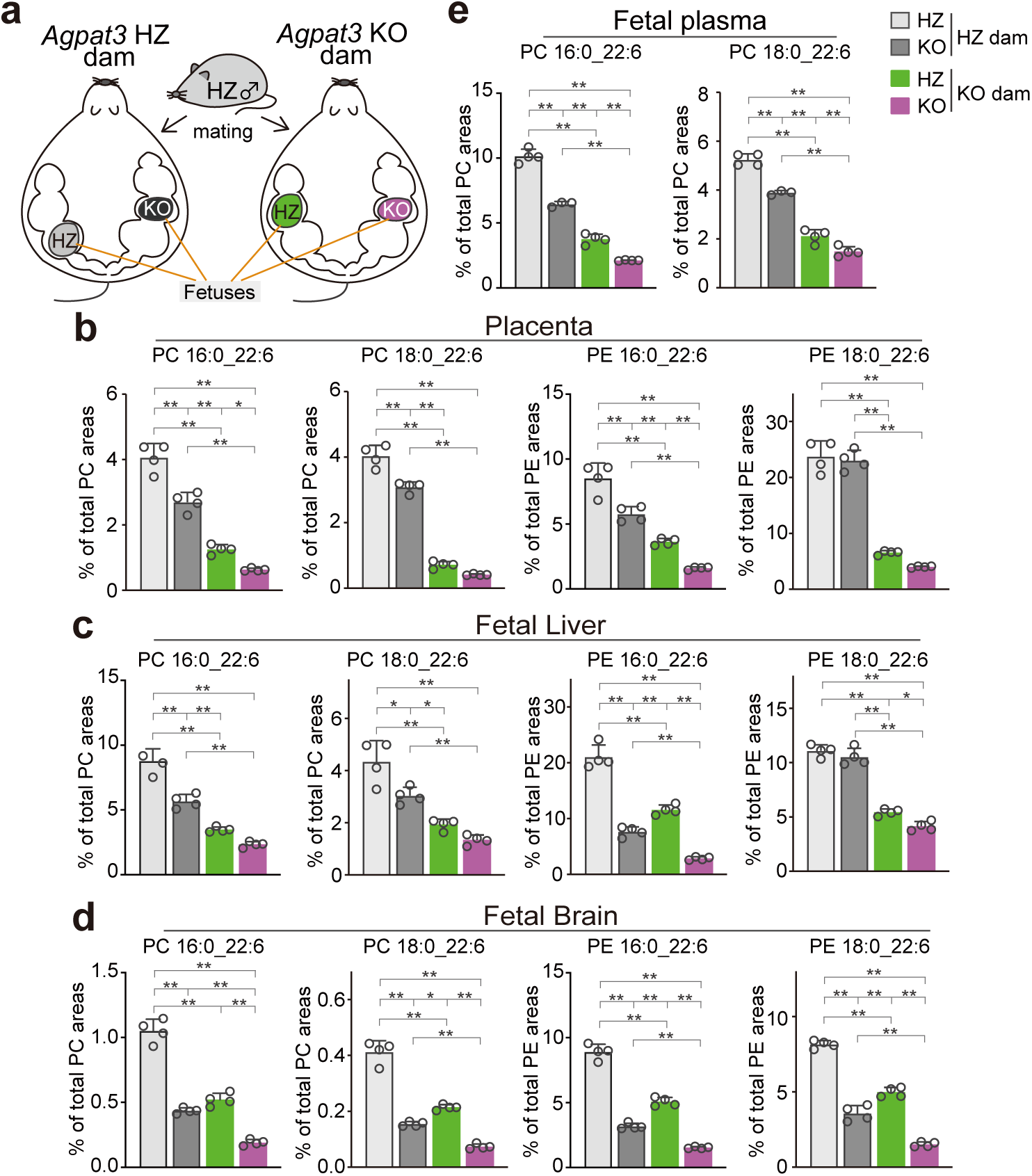
Impact of maternal–fetal *Agpat3* genotypes on fetal tissue docosahexaenoic acid-containing phospholipid (DHA-PL) levels. (**a**) Schematic diagram of the procedure to generate fetuses with different maternal–fetal *Agpat3* genotype combinations. (**b**–**e**) Relative abundances of DHA-containing phosphatidylcholine (DHA-PC; PC 16:0_22:6 and PC 18:0_22:6) and phosphatidylethanolamine (DHA-PE; PE 16:0_22:6 and PE 18:0_22:6) in the placenta (**b**), fetal liver (**c**), fetal brain (**d**), and fetal plasma (**e**) of embryonic day 18 mice. Light gray (*Agpat3* HZ) and dark gray (*Agpat3* KO) indicate fetuses of *Agpat3*-HZ dams. Green (*Agpat3* HZ) and magenta (*Agpat3* KO) indicate fetuses of *Agpat3*-KO dams. Data are presented as means + SD (n = 4 for each group). Statistical significance was determined using Bonferroni’s multiple comparisons (\**P* < 0.05, \*\**P* < 0.01).

Subsequently, we investigated the effect of DHA-PL deficiency in dams on the DHA-PL levels in their offspring during neonatal development. We analyzed DHA-PC and -PE levels in the neonatal liver and brain on postnatal day 1 (P1), P8, and P28 (Fig. 4a). On P1, the maternal *Agpat3* genotype evidently affected the DHA-PL levels in *Agpat3*-HZ and -KO mice, as observed in the fetal stage (Fig. 4b, c). However, the influence of the maternal *Agpat3* genotype on the DHA-PL levels in the neonatal tissues was completely abolished after P8, and DHA-PL levels entirely depended on the fetal *Agpat3* genotype (Fig. 4b, c).

**Fig. 4.**
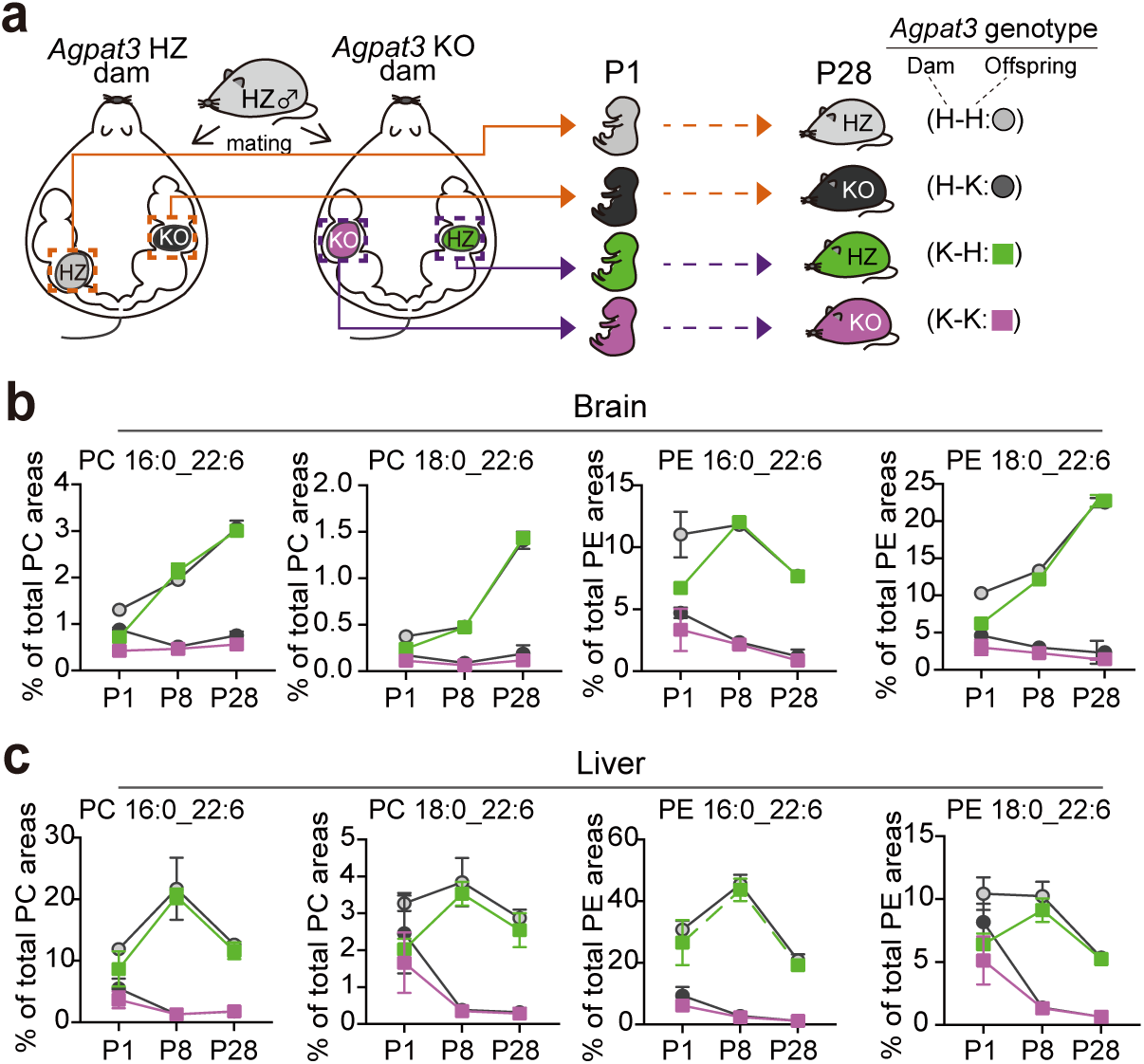
Postnatal temporal changes in docosahexaenoic acid (DHA)-containing phospholipid levels in neonatal mouse tissues. Schematic diagram of the procedure to generate neonates with different maternal–fetal *Agpat3* genotype combinations. (**a**–**c**) Light gray (*Agpat3* HZ, represented as H-H) and dark gray (*Agpat3* KO, represented as H-K) indicate offspring of *Agpat3*-HZ dams. Green (*Agpat3* HZ, represented as K-H) and magenta (*Agpat3* KO, represented as K-K) indicate offspring of *Agpat3*-KO dams. (**b**, **c**) Line graphs depicting the relative abundances of DHA-containing phosphatidylcholine (DHA-PC) and DHA-phosphatidylethanolamine (DHA-PE) in the brain (b) and liver (**c**) of postnatal day 1 (P1), P8, and P28 mice. Data are presented as means ± SD (n = 4 for each group).

The source of maternal DHA supply to the offspring changes from maternal blood to breast milk fat, which is composed of over 95% TGs ^15^. The abolished effect of the maternal *Agpat3* genotype on fetal tissue DHA-PL levels (Fig. 4b, c) may be explained by a switch in maternal lipid sources from maternal blood to breast milk after birth considering that the *Agpat3* genotype affects the levels of DHA-PL but not DHA-TG. Concordantly, gas chromatography (GC) of breast milk of *Agpat3*-HZ and -KO dams revealed no significant differences in the total fatty acid composition, including DHA (Supplementary Fig. 4). These results suggest that the source of DHA for infants is most likely maternal DHA-TGs, rather than DHA-PLs.

### Docosahexaenoic acid-PL deficiency in the neonatal stage (but not the fetal stage) leads to reductions in body weight and brain size

Our lipidomic data revealed that the *Agpat3*-KO mouse model is suitable to separately examine the impact of DHA-PL deficiency in the fetal and postnatal periods (Fig. 5a). We compared the body weight and brain size of offspring among four combinations of dam–offspring *Agpat3* genotypes to assess the effect of DHA-PL deficiency in each stage on offspring growth (Fig. 5a). Despite different DHA-PL levels in the fetal stages, no significant difference in body weight was observed among the dam–offspring *Agpat3* genotypes from embryonic day 18 to P8 (Fig. 5b). A slightly decreasing trend in brain weight was observed on P8 (Fig. 5c). However, *Agpat3*-KO mice had reduced body and brain weight, irrespective of the genotype of the dams on P28 (Fig. 5b, c). Collectively, these results indicate that the deficiency of DHA-PLs hinders brain size and body weight gains in the postnatal stage but not the fetal stage.

**Fig. 5.**
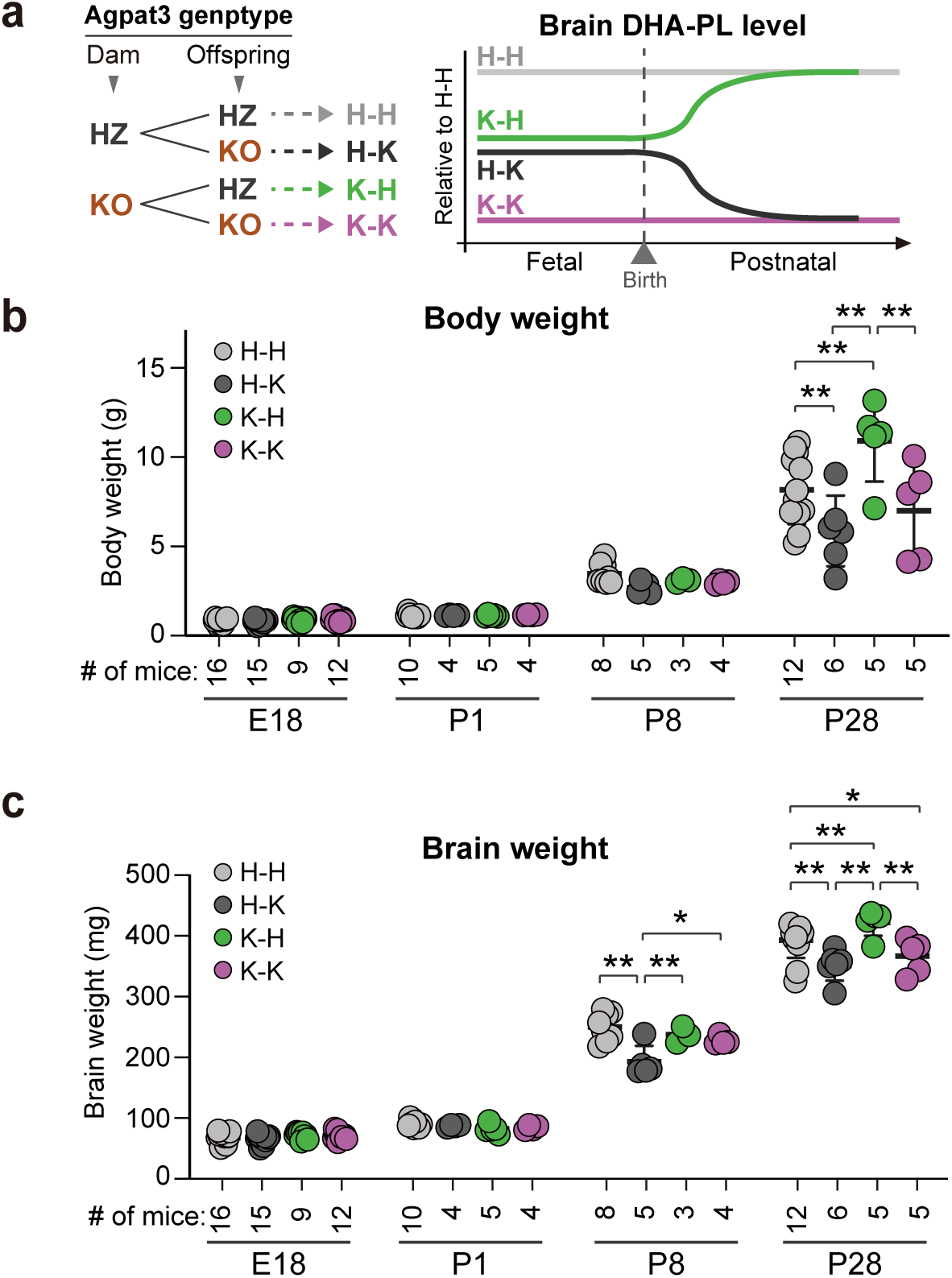
Impact of *Agpat3* deficiency on body weight and brain weight from the fetal to the weaning stages in mice. (**a**) Summary of the effects of maternal–offspring *Agpat3* genotypes on brain docosahexaenoic acid-containing phospholipid (DHA-PL) levels in the offspring. (**a**–**c**) Light gray (*Agpat3* HZ, represented as H-H) and dark gray (*Agpat3* KO, represented as H-K) indicate offspring of *Agpat3*-HZ dams. Green (*Agpat3* HZ, represented as K-H) and magenta (*Agpat3* KO, represented as K-K) indicate offspring of *Agpat3*-KO dams. (**b**, **c**) Body weight (**b**) and brain weight (**c**) on embryonic day 18 (E18), postnatal day 1 (P1), P8, and P28 in mice with different dam–fetal *Agpat3* genotype combinations. Each dot represents an individual mouse, and mean ± SD is indicated. The numbers of mice are indicated below each dataset. Statistical significance was determined using Bonferroni’s multiple comparisons (\**P* < 0.05, \*\**P* < 0.01).

### Potential impact of prenatal DHA-PL deficiency on prospective brain functions in mice

Various clinical interventions and animal models suggest that DHA deficiency causes brain structural abnormalities and behavioral disorders ^28, 29^. We compared the motor function of *Agpat3*-WT, -HZ, and -KO mice nursed by a surrogate (ICR) dam to determine whether *Agpat3* deletion affects neurological functions in mice. Compared with *Agpat3*-WT or -HZ mice, *Agpat3*-KO mice spent significantly less time on the rotating rod (Fig. 6a). Furthermore, compared with *Agpat3*-WT or -HZ mice, *Agpat3*-KO mice exhibited significantly increased hindlimb clasp severity (Fig. 6b). These results suggest that DHA-PL deficiency leads to severe motor dysfunction, consistent with findings in other mouse models of DHA deficiency ^29, 30, 31, 32, 33^.

**Fig. 6.**
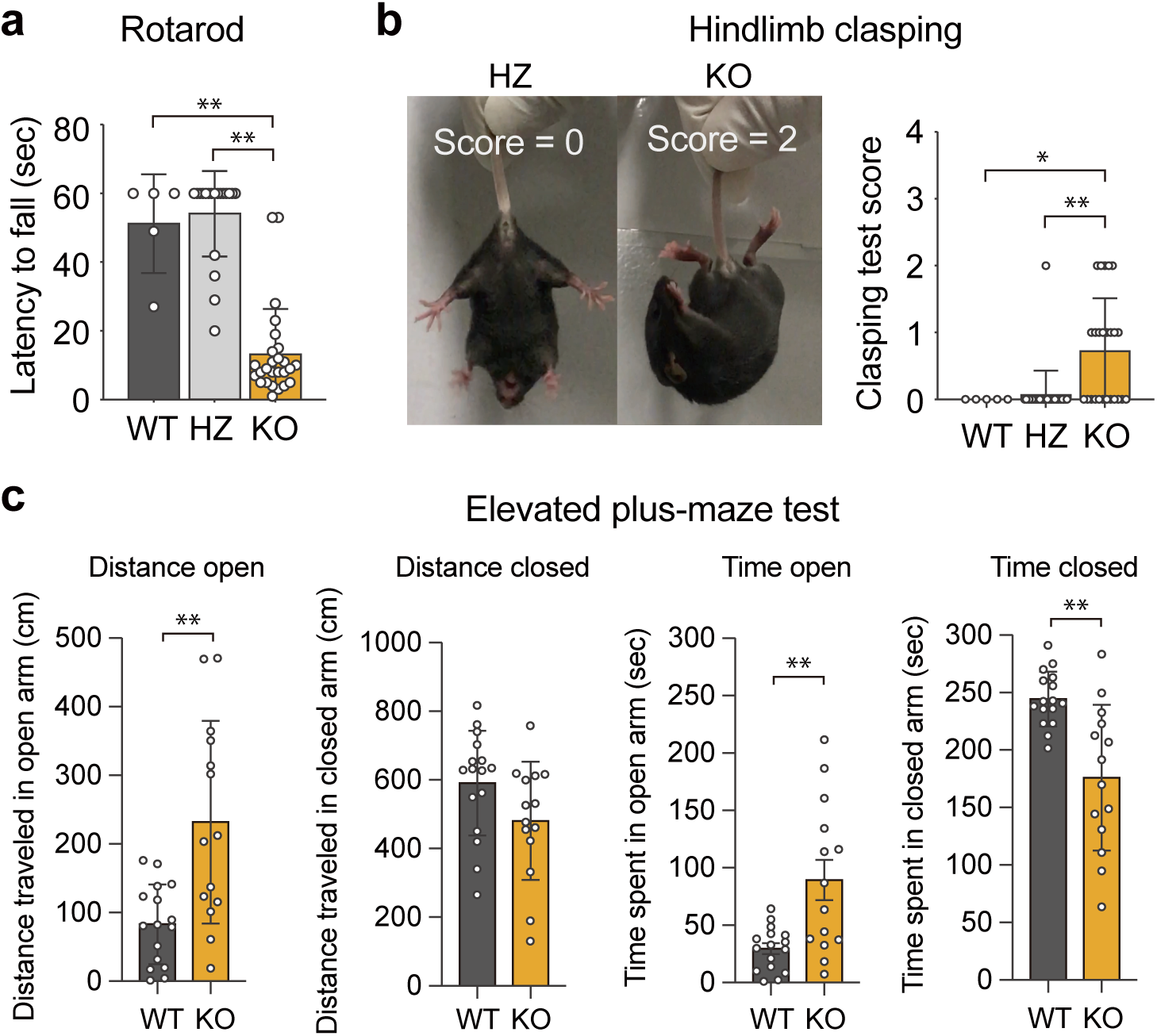
Behavioral phenotypes of *Agpat3*-KO mice. (**a**–**c**) Results of hindlimb clasping (**a**), rotarod (**b**), and elevated plus-maze (EPM) (**c**) tests. (**a**) The hindlimb clasping test was performed using 4-week-old mice. The hindlimb clasping reflex was scored on a scale of 0–4, as described in the Methods section. Representative images (left) of the hindlimb clasping phenotypes. *Agpat3*-HZ and -KO mice with scores of 0 and 2, respectively, are presented. The bar graphs (right) depict the hindlimb clasping scores of *Agpat3*-WT, -HZ, and -KO mice (n = 5, 30, and 25, respectively). (**b**) The rotarod test was performed using 10-week-old mice. Falling latency of *Agpat3*-WT, -HZ, and -KO mice (n = 5, 19, and 26, respectively) on a rotating rod at 10 rpm (60-s cutoff). (**a**, **b**) Data are presented as means ± SD. Statistical significance was determined using Bonferroni’s multiple comparisons (\**P* < 0.05, \*\**P* < 0.01). (**c**) Results of EPM tests of 8-week-old *Agpat3*-WT and -KO mice. Time and distance in the open and closed arms are presented. Each dot represents an individual mouse (n = 16 for WT and n = 14 for KO). Data are presented as means ± SD. Statistical significance was determined using an unpaired *t*-test (\**P* < 0.01).

In the elevated plus-maze (EPM) test, the time and distance traveled in the open arms were significantly longer for *Agpat3*-KO mice than for *Agpat3*-WT mice (Fig. 6c), suggesting that DHA-PL deficiency is associated with reduced anxiety-like behavior. Despite these behavioral abnormalities, no apparent histological abnormalities were observed in *Agpat3*-KO mouse brains (Supplementary Fig. 5). This differs from major facilitator superfamily domain-containing 2a (*Mfsd2a*)-KO mice, another model of DHA deficiency ^33^.

DHA is considered a critical nutrient for brain development in fetuses and neonates. However, the impact of DHA deficiency in the fetal brain on prospective brain functions remains obscure. Therefore, we compared the motor functions of *Agpat3*-KO mice with different brain DHA-PL levels only in the fetal stages (*Agpat3*-HZ mice born from either *Agpat3*-WT or -KO dams) (Fig. 7a). *Agpat3*-HZ offspring from *Agpat3*-KO dams displayed a performance comparable with that of offspring from *Agpat3*-WT dams in rotarod and hindlimb clasping tests, suggesting that DHA-PL deficiency in the fetal brain does not affect prospective motor functions (Supplementary Fig. 6). In contrast, the EPM test revealed that the maternal genotype did affect the behavior of their offspring (Fig. 7b, c). Male *Agpat3*-HZ mice born from *Agpat3*-KO dams traveled significantly longer distances (*P* = 0.020) and tended to spend more time (*P* = 0.056) in the open arms than those delivered by *Agpat3*-WT dams (Fig. 7b). Female *Agpat3*-HZ offspring from *Agpat3*-KO dams tended to travel longer distances (*P* = 0.070) and spent significantly more time (*P* = 0.030) in the open arms than offspring from *Agpat3*-WT dams (Fig. 7c). Together, these findings suggest that fetal stage-specific brain DHA-PL deficiency can lead to a reduction in anxiety-like behavior during adulthood. Therefore, these results support the notion that DHA deficiency in the fetal stage can influence prospective neuropsychiatric function.

**Fig. 7.**
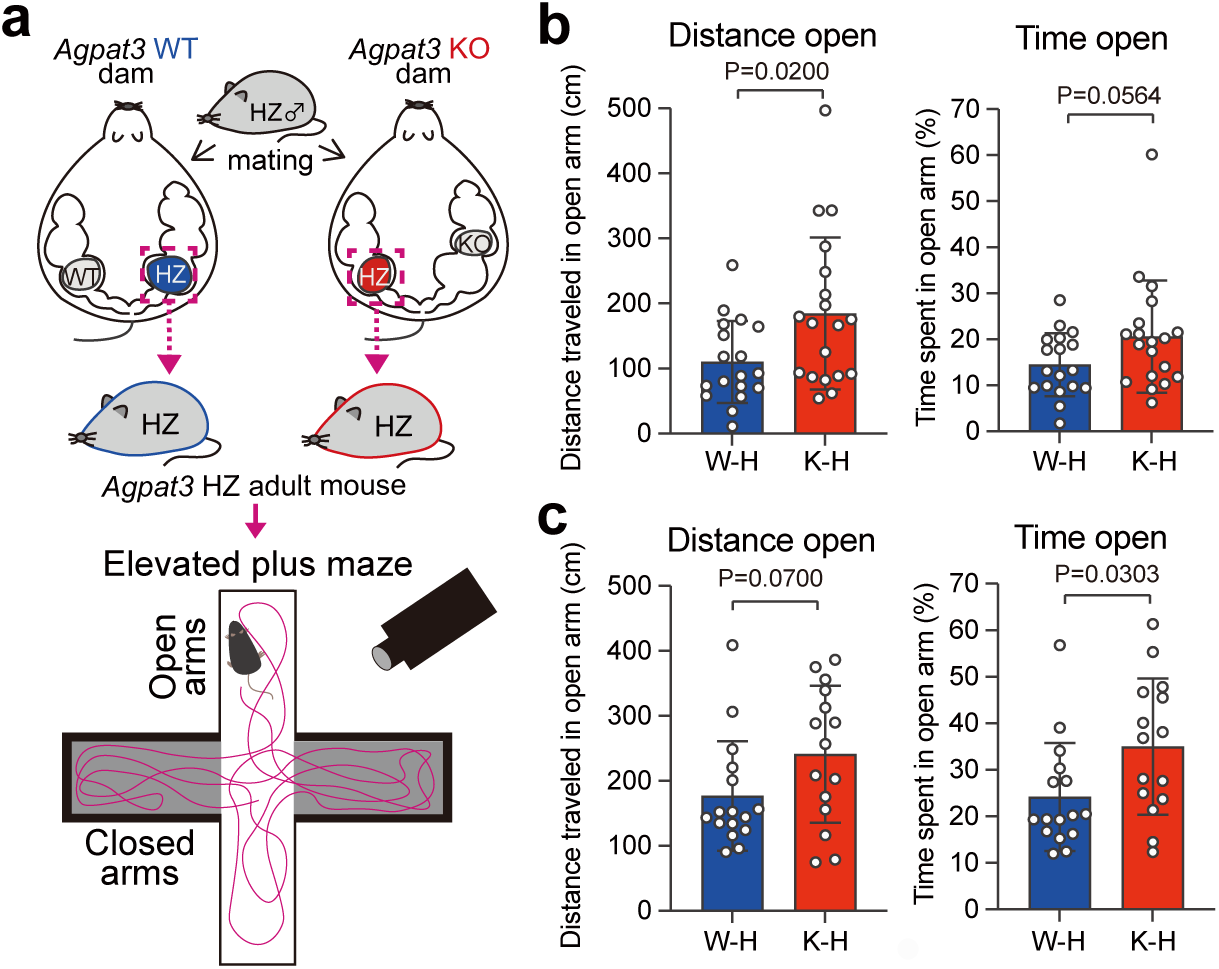
Impact of fetal stage-specific docosahexaenoic acid-containing phospholipid (DHA-PL) deficiency on anxiety-or fear-related behavior of adult mice. (a) Schematic representation of the behavioral assessment of *Agpat3*-HZ mice born from *Agpat3*-WT or -KO dams using the elevated plus-maze (EPM) test. (**b**, **c**) Results of EPM tests of 14–16-week-old male (**b**) and female (**c**) *Agpat3*-HZ mice born from *Agpat3*-WT or -KO dams. Distance (left) and time (right) in the open arm are shown. W-H and K-H indicate the maternal–fetal *Agpat3* genotype (W, H, and K indicate WT, HZ, and KO, respectively). Each dot represents an individual mouse (n = 17 for male W-H, n = 19 for male K-H, n = 16 for female W-H, and n = 15 for female K-H) and mean ± SD is presented. Presented *P*-values were determined using an unpaired *t*-test

## Discussion

The limited understanding of the complex mechanisms underlying maternal–fetal DHA transfer hinders our ability to discern the specific roles of DHA-PLs and DHA-TGs in maternal blood, thereby limiting targeted interventions to address omega-3 fatty acid deficiencies during critical developmental periods. This study demonstrated that the loss of DHA-PL synthesis in dams affected the DHA supply to the offspring during the fetal stages but not in infancy. Although fetal stage-specific DHA-PL deficiency had no evident impact on postnatal DHA-PL levels, brain gross histology, and growth in the offspring, it influenced their prospective neurological phenotypes.

Our findings suggest that DHA-PLs are a vital source of fetal DHA supply among the various types of DHA-containing lipids in maternal blood. Concordantly, a recent study using delta-like homolog 1 gene-KO mice suggested a major contribution of PLs in the maternal blood to dam–the fetal transfer of polyunsaturated fatty acids, including DHA ^34^. Therefore, the results of this study and previous reports ^16, 17, 18^ show that the increase in DHA-PLs in maternal blood during pregnancy may contribute to fulfilling the increased DHA requirement in fetal tissues (particularly in the brain) in the late stage of pregnancy ^35^. Hepatocytes are likely to be involved in the increase in DHA-PLs in the blood during pregnancy considering that circulatory DHA-PL levels largely depend on DHA synthesis by hepatic AGPAT3 ^23^. Concurrently, enhanced hepatic expression of enzymes involved in DHA and DHA-PL synthesis is demonstrated in pregnant rodents ^16, 34^.

The current findings prompt the question regarding the manner in which maternal circulatory DHA-PLs function as a DHA source for the fetus. Besides the most commonly discussed possibility that DHA in DHA-PLs is transported after cleavage by endothelial lipase ^36^, several other pathways may be involved in the transplacental transfer of DHA in PLs to the fetus. Initially, DHA-PLs in the blood can be converted to DHA-lysoPLs that may be transported to the fetus via MFSD2A, a lysoPL transporter critical for DHA transport across the blood–brain barrier. Concurrently, human placental syncytiotrophoblasts highly express MFSD2A ^37^. Furthermore, the *MFSD2A* expression level in the placenta correlates with the cord blood DHA-PL concentration in humans ^38^, further confirming the potential involvement of MFSD2A in transplacental DHA transfer. Second, DHA-PLs within lipoproteins may be directly incorporated into syncytiotrophoblasts via scavenger receptors such as scavenger receptor class B type 1 (SR-BI). In mice, SR-BI is expressed at maternal–fetal interfaces, such as the placenta and yolk sac, and its deficiency causes fetal developmental defects, including anencephaly ^39^. In this scenario, some of the DHA-PLs incorporated within lipoproteins may be directly transferred into fetal circulation via transcytosis across the placenta by bypassing fatty acid remodeling in the syncytiotrophoblasts. However, we lack evidence supporting the contributions of MFSD2A and SR-BI to transplacental DHA transport.

The impact of perinatal DHA deficiency on brain expansion differs among experimental DHA deficiency models; therefore, it remains controversial. Rats fed an omega-3-deficient diet for several generations, and mice deficient in acyl-CoA synthetase long-chain family member 6 (*Acsl6*) and fatty acid-binding protein 5 (*Fabp5*), revealed no difference in brain weight compared with that in control animals ^31, 32, 40, 41^. However, deletion or dysfunction of the gene encoding MFSD2A causes microcephaly in mice and humans ^42, 43, 44^. Our findings suggest that the loss of *Agpat3* in mice led to a decrease in brain weight at 4 weeks of age. However, histological abnormalities observed in *Mfsd2a*-KO mice (such as decreased numbers of Purkinje cells and hippocampal neurons) were not detected in the *Agpat3*-KO mouse brain^33^. This result is consistent with a recent study showing no prominent abnormalities in MRI and CT scans in patients harboring a homozygote nonsense variant of *AGPAT3* (c.747C>A), which generates a truncated (p. Tyr249Ter) and unstable form of AGPAT3 ^45^.

The requirement of DHA for normal brain function was demonstrated by animal behavioral tests of omega-3 fatty acid-deficient diet-fed animals and KO mouse models with reduced brain DHA levels ^29, 30, 31, 32, 33^. The neuropsychiatric abnormalities observed in *Agpat3*-KO mice (Fig. 6) demonstrate the importance of DHA in brain function. The importance of AGPAT3 in brain function is further supported by a recent study that identified a homozygote *AGPAT3* mutation in individuals with symptoms of severe intellectual disability and retinitis pigmentosa ^45^. The aggressive behavioral phenotype of patients harboring this *AGPAT3* mutation ^45^ can be explained by the reduced anxiety-like behavior of *Agpat3*-KO mice in the EPM test. However, rodent model animals on omega-3 fatty acid-deficient diets have yielded inconsistent EPM test results since EPM test results are affected by experimental conditions, such as maze height and brightness ^10^. Therefore, further studies are required to reveal the detailed effects of DHA-PL deficiency on brain function.

Our results further suggest that lifelong DHA-PL deficiency together with fetal stage-specific DHA-PL deficiency can affect adult brain functions. However, the mechanisms by which DHA-PL deficiency affects the behavior of adult mice during fetal stages remains unclear. DHA-PL deficiency in the fetal stage (approximately 50% reduction) may have caused an improper neuronal distribution that can affect neurological functions in adulthood since short hairpin RNA-mediated suppression of *AGPAT3* (approximately 50% reduction in *AGPAT3* mRNA in the brain) impairs neuronal migration in fetal mice ^45^. Furthermore, epigenetic changes in the developing fetal brain caused by DHA deficiency likely influence prospective brain functions. Notably, omega-3 fatty acid levels affect epigenetic status, such as DNA methylation and histone modifications ^46, 47^. Comparative spatial transcriptomics and neuronal epigenomic profiles of brains of *Agpat3*-HZ mice born from *Agpat3*-WT and -KO dams would deepen our understanding of the impact of DHA deficiency in the fetal stage on brain functions in adults.

However, our study has a few limitations. Although we used a mouse model to investigate maternal–fetal DHA transport and the effects of DHA deficiency on brain development, there are non-negligible differences in perinatal brain development between rodents and humans. In mice, the peak of brain expansion and neuronal differentiation (termed “brain growth spurt”) occurs from one week after birth to weaning ^48^. In humans, the brain growth spurt occurs between 36–40 weeks of gestation ^14^. Furthermore, neurogenesis continues after birth in mice, whereas it finishes before birth in humans ^49^. Therefore, the impact of maternal–fetal DHA supply on brain development and prospective brain functions may be more pronounced in humans than in mice.

Despite these limitations, our findings highlight the critical role of maternal DHA-PL synthesis in fetal DHA supply and neurological outcomes, emphasizing DHA-PLs as a key source to meet increased DHA requirements in fetal tissues, particularly the brain. Neurological abnormalities in *Agpat3*-KO mice highlight the importance of DHA incorporation into PLs for normal brain function, suggesting implications for human health. Overall, our findings provide novel insights into maternal-fetal DHA transport, laying the groundwork for potential therapeutic strategies supporting perinatal brain development in the offspring.

## Methods

### Animals

Mice were housed at 23 ± 2°C and 40–60% humidity with a 12-hour light/dark cycle under a specific pathogen-free (SPF) condition and provided *ad libitum* access to a standard diet (CE-2, CLEA, Japan) and drinking water. C57BL/6J mice were purchased from CLEA Japan. Pregnancy was confirmed by the presence of a vaginal plug, and the day on which the vaginal plug was confirmed was day 0 of pregnancy. All animal experiments were approved by the President of NCGM, following consideration by the Institutional Animal Care and Use Committee of NCGM (approval ID: 17054, 17085, 17093, 17100, 18046, 18084, 18096, 18119, 19039, 19093, 19112, 20035, 20045, 20093, 20106, 21023, 21039, 22052, 2023-A065) and were performed in accordance with institutional procedures, national guidelines, and the relevant national laws on the protection of animals.

### *Agpat3* KO mouse

*Agpat3* KO mice (Agpat3tm1(EUCOMM)Wtsi) were purchased from the European Conditional Mouse Mutagenesis Project. *Agpat3* was deleted by the insertion of a LacZ cassette with a splice acceptor and a poly A signal between exons 3 and 4 of the *Agpat3* gene, which is transcribed as a fusion of exons 3 and the *LacZ* gene. The genotype of each mouse was confirmed by PCR using three primers (WT forward, accttatgaaggtgaccatgtggag; KO forward, cgtcgagaagttcctattccgaagt; Common reverse, gtgtcctgaatgaccaggaagagaa) as previously reported ^26^. The expected product sizes are 381 bp for the WT allele and 244 bp for the KO allele.

### Tissue and blood collection

Mouse tissue and blood samples were collected under anesthesia with sodium pentobarbital. Tissues were frozen in liquid nitrogen immediately after sampling and stored at −80°C. Adult mouse blood collected from the abdominal aorta was gently transferred into a Fuchigami separation tube (Matsume Seisakusho, Japan) containing a coagulation-accelerating separator, allowed to settle at room temperature for 30 min, and centrifuged at 1000 x *g* for 5 min at 4°C. The supernatant representing the serum was frozen in liquid nitrogen stored at −80°C. Fetal blood was collected using a heparin-coated capillary tube. Collected blood was kept on ice and centrifuged at 14,000 x *g* 5 min at 4°C. The supernatant was collected as fetal plasma, frozen in liquid nitrogen, and stored at −80°C.

### Lipid extraction

Frozen tissues were ground to a powder using a cryo-crusher (Automill; Tokken inc., Japan). Methanol (for PL analyses) or isopropyl alcohol (for neutral lipid analyses) was added to the tissue powder, and lipids were extracted with gentle agitation by a rotator of extraction tubes for 1 h at 4°C with rotation. Subsequently, the tubes were centrifuged at 20,000 × *g* for 10 min at 4°C and the supernatants were used for lipid analyses. Frozen tissues were used for lipid and RT-qPCR analyses. Tissues were homogenized using Handy Micro Homogenizer (NS310E2, Microtec Co. Ltd., Japan) with 40 µL / mg tissue of RLT buffer (RNeasy Mini Kit, Qiagen, USA), which contained 10 µL / mL of β-mercaptoethanol (β-ME). Subsequently, the lipids were extracted from the tissue homogenate by the Bligh and Dyer method ^50^. The extracted lipid fractions were dried by centrifugal evaporator (TAITEC, Japan). In the case of blood (serum and plasma), samples were dissolved in methanol (for PL analyses) or isopropyl alcohol (for neutral lipid analyses) and mixed using a vortex mixer for 1 min. Subsequently, the samples were centrifuged at 16,000 x *g* for 5 min at room temperature. The supernatant was used for the lipid analyses.

### Annotation of lipids

The nomenclature of lipids followed recently proposed rules ^51^. Fatty acid species were indicated as “FA AA:B (n-Y)”; “FA” represents fatty acid, “AA” represents a number of carbons, “B” represents number of double bonds, and “Y” represents the carbon number that possesses the first double bond from its omega-end. Fatty acid species in LPC and CE were represented as X AA:B (“X” indicates the lipid class). Fatty acid composition of PC, PE, and TG were denoted as X CC:D (“CC” and “D” represent the sum of carbon number and double bonds of fatty acids within the lipid, respectively). In the case of DHA-PC and -PE, these PLs were indicated as “X AA:B_22:6”. DHA-TG or TG CC:D (22:6) indicates the presence of at least one DHA moiety in its structure. We did not determine the *sn* position that each fatty acid harbors.

### Liquid chromatography/tandem mass spectrometry analyses of PL fatty acid compositions

Fatty acid composition of PC, PE and LPC was analyzed by multiple-reaction monitoring (MRM) using LC-MS/MS (LCMS-8050, Shimadzu Corp., Japan). Extracted lipid samples were separated on a column [Acquity UPLC BEH C8 column (1.7 µm, 2.1 × 100 mm, Waters, USA)] heated to 47°C using ternary mobile phase gradient (mobile phase A: 5 mM ammonium bicarbonate; mobile phase B: acetonitrile; mobile phase C: isopropyl alcohol) at a flow rate of 0.35 mL/min. The mobile phase gradient was set as follows: [time (% A/% B/% C): 0 min (50/45/5), 10 min (20/75/5), 20 min (20/50/30), 27.5 min (5/5/90), 28.5 min (5/5/90), 28.6 min (50/45/5)]. The MRM settings for measuring each PL were set as follows: (Q1, Q3): PC and LPC ([M + H]^+^, 184), PE ([M + H]^+^, neutral loss of 141). The peak annotation, deisotoping, and calculation of the amount (area under the curve) of each peak were performed using TRACES, a software for MRM-based LC-MS analyses ^52^. The peaks of DHA-PLs were identified by the column retention time as previously described ^26^.

### Liquid chromatography/tandem mass spectrometry analyses of fatty acid compositions in TG and CE

The lipid composition of TG and CE was also analyzed by MRM using LC-MS/MS (LCMS-8060, Shimadzu Corp., Japan). Samples for lipid analysis were separated on a column [Acquity UPLC BEH C8 column (1.7 µm. 2.1 x 100 mm, Waters, USA)] heated to 47°C using mobile phase A: 10 mM ammonium acetate / 0.1% formic acid; mobile phase B: ammonium acetate / 0.1% formic acid / 99% methanol; mobile phase C: isopropyl alcohol at a flow rate of 0.2 mL/min. The gradient of the mobile layer used for the separation was set as follows: gradient analysis [time (% A/% B/% C): 0 min (30/20/50), 12 min (5/45/50), 15 min (10/10/80), 19.1 min (30/20/50)]. The MRMs for the measurement of each lipid class was set as follows. (Q1, Q3): TG ([M + NH_4_]^+^, neutral loss of fatty acids and ammonium), CE ([M + NH_4_]^+^, 369.4). The peak annotation and calculation of amount (area under the curve) of each peak were performed using TRACES ^52^.

### Gas chromatography-flame ionization detector (GC-FID)

Solidified contents were collected from the gastric of 8-day-old neonatal mice as breast milk. Breast milk samples were frozen in liquid nitrogen and stored at −80°C until use. After adding tricosanoic acid (FA 23:0) (Sigma-Aldrich, USA) as an internal standard, total fatty acids were methyl esterified using a Fatty acid methylation kit (Nacalai Tesque, Japan) and purified using a fatty acid methyl ester purification kit (Nacalai Tesque). The resulting solution was dried in a centrifugal evaporator, re-dissolved in dichloromethane, and used for GC-FID analyses. Docosatetraenoic acid (DTA) methyl ester, docosapentaenoic acid (DPA) methyl ester, and Supelco 37-component FAME Mix (Merck, Germany) were used to identify and quantify each fatty acid. A GC system equipped with a hydrogen flame ionization detector (GC-2010 Plus, Shimadzu, Japan) was used for GC-FID analysis. The injector and detector temperatures were set to 240 and 250°C, respectively, and the helium carrier gas flow velocity was 45 cm / s. The FAME species were separated by a FAMEWAX column (12497, Restek, USA) with the following temperature gradient (rate of temperature increase (°C / min)); starting at 140°C, and increased to 200°C (11°C / min), 225°C (3°C / min), 240°C (20°C / min), and 240°C for 5 min. Fatty acid concentrations were determined using GCsolution (Shimadzu Corp., Japan), a dedicated analysis software. Fatty acid concentrations in each sample were normalized by tricosanoic acid (FA 23:0) ^53^.

### Serum TG amounts

Total TG in serum was determined using the LabAssay Triglyceride Kit (Wako, Japan) according to the accompanying manual.

### Quantitative RT-PCR

Freshly frozen tissues were homogenized as described above (see “Lipid extraction”). Total RNA was extracted from the supernatant (12,000 x *g* for 10 min at 4°C) of centrifuged tissue homogenate using an RNeasy Mini Kit (Qiagen, USA). The extracted total RNA was used for single-strand cDNA synthesis. Single-stranded cDNA was synthesized using the reverse transcriptase Superscript III and random primers (Thermo Fisher Scientific). Quantitative PCR was performed with 2 µL of cDNA as a template in a total volume of 20 µL containing LightCycler FastStart DNA MasterPlus SYBR Green I (Roche, USA) and 0.5 µM primer. The following primer sequences were used (5’-3’): (Gene name: Forward, Reverse) *Fads1*: GAAGAAGCACATGCCATACAACC, TCCGCTGAACCACAAAATAGAAA; *Fads2*: GCCTGGTTCATCCTCTCGTACTT, GAAAGGTGGCCATAGTCATGTTG; *Elovl2:* CCTGCTCTCGATATGGCTGG, AAGAAGTGTGATTGCGAGGTTAT; *Elovl5*: ATGGAACATTTCGATGCGTCA, GTCCCAGCCATACAATGAGTAAG; and *Rplp0*: CTGAGATTCGGGATATGCTGTTG, AAAGCCTGGAAGAAGGAGGTCTT. The mRNA expression levels were calculated by the ΔΔCt method. *Rplp0* was used for the internal control.

### Histological analysis

Nissl-stained brain sections were prepared at Genostaff (Tokyo, Japan). Briefly, mice were deeply anesthetized and transcardially perfused with PBS for 5 min and 4% paraformaldehyde (PFA) for 60 min at room temperature. Brains were dissected and post-fixed in 4% PFA overnight at room temperature followed by embedding in paraffin blocks. Sagittal brain sections (6 µm) were subjected to Nissl staining. Stained slides were digitally scanned using a NanozoomerS210 (Hamamatsu Photonics, Hamamatsu, Japan). The NDP.view 2 software (Hamamatsu Photonics) was used for histological assessment including counting the number of neurons in the hippocampal regions CA1 and CA3, and Purkinje cells in cerebellum, and to measure the length of these regions as a correcting factor.

### Rotarod test

The apparatus consisted of five rotating rods separated by walls. Ten-week-old mice were set on the rotating rod at 2 rpm, and the rotation was accelerated to 10 rpm. The time at which the rotational speed was set to 10 rpm was set to 0, and the time until the mouse fell off the rotating rod (60 s cut-off) was measured. The experimenter was blind to mouse genotype throughout the testing.

### Hindlimb clasping test

Four-week-old mice were suspended by the middle of the tail and their behaviors were recorded for 60 s. Hindlimb clasping was scored from 0 to 4 based on severity: 0 = both hindlimbs were splayed outward away from the abdomen with splayed toes, 1 = one hindlimb was retracted toward the abdomen, 2 = both hindlimbs were retracted toward the abdomen, 3 = both hindlimbs were retracted and touched the abdomen, 4 = both hindlimbs were fully clasped and touched the abdomen. Scoring was performed by an individual blind to the genotype of the mice.

### Elevated plus-maze

The test apparatus consisted of two open arms and closed arms (25 x 5 cm) that extended from a central platform (5 x 5 cm). The height of the closed arm walls was 15 cm, and the maze was elevated 70 cm above the floor. Fourteen-sixteen-(Fig. 6) and eight-week-old mice (Supplementary Fig. 5c) were transferred to the testing room 1 h prior to testing for acclimation to the test environment. The mice were placed on a central platform and allowed to investigate the maze for 5 min with the activity recorded and tracked by LimeLight software (Actimetrics). Data recorded included distance traveled in each region and percent time spent in the open arms of the maze. The tests were conducted between 9:00 and 17:00 h. The observer was blind to genotypes and experimental conditions.

### Statistical analyses

Unpaired *t*-tests were used to compare two groups. Bonferroni’s multiple comparison tests were used to compare three or more groups. All statistical analyses were performed using GraphPad Prism 9 (version 9.5.1) software.

## Acknowledgments

This work was supported by the Japan Agency for Medical Research and Development (AMED)-CREST (grant number JP22gm0910011 to H.S.), the AMED Program for Basic and Clinical Research on Hepatitis (grant number 23fk0210091 to H.S.), the AMED-PRIME (grant number 22gm6310018 and 23gm6710020 to K.Y.), Japan Society for the Promotion of Science [grant number 20K08900 and 23K18103 to K.Y.], the National Center for Global Health and Medicine, Intramural Research Fund (29-1033 to D.H., 22T001 to H.S., and 19A1023 and 22A1012 to K.Y.), the Japan Health Research Promotion Bureau Research Fund (JH2022-B-03 to H.S.), the TERUMO LIFE SCIENCE FOUNDATION (22-ΙΙΙ4054 to H.S.), Takeda Science Foundation (to T.S. and K.Y.), Astellas Foundation for Research on Metabolic Disorders (to K.Y.), the Cell Science Research Foundation (to K.Y.), the Mitsubishi Foundation (to K.Y.), Japan Foundation for Applied Enzymology (to K.Y.), and the Uehara Memorial Foundation (to K.Y.).

We thank Takehiko Sasaki and Junko Sasaki (Tokyo Medical and Dental University, Japan) for providing *Agpat3* KO mouse. We thank Yukito Ishizaka (Department of Intractable Diseases, National Center for Global Health and Medicine, Japan) for technical assistance with the rotarod test.

## Author contributions

A.K., D.H., K.Nag., T.N., and K.Y. conceptualized the study.

A.K., D.H., K.Nag., F.H., and K.Y. performed the experiments and data analyses.

K.Nak. and T.O. performed IVF.

A.K., D.H., and K.Y wrote the manuscript.

K.Nag, F.H., H.S., T.S., and T.N. reviewed and edited the manuscript.

## Materials and Correspondence

Please address correspondence to Daisuke Hishikawa and Keisuke Yanagida

---

**Supplementary Fig. 1.**
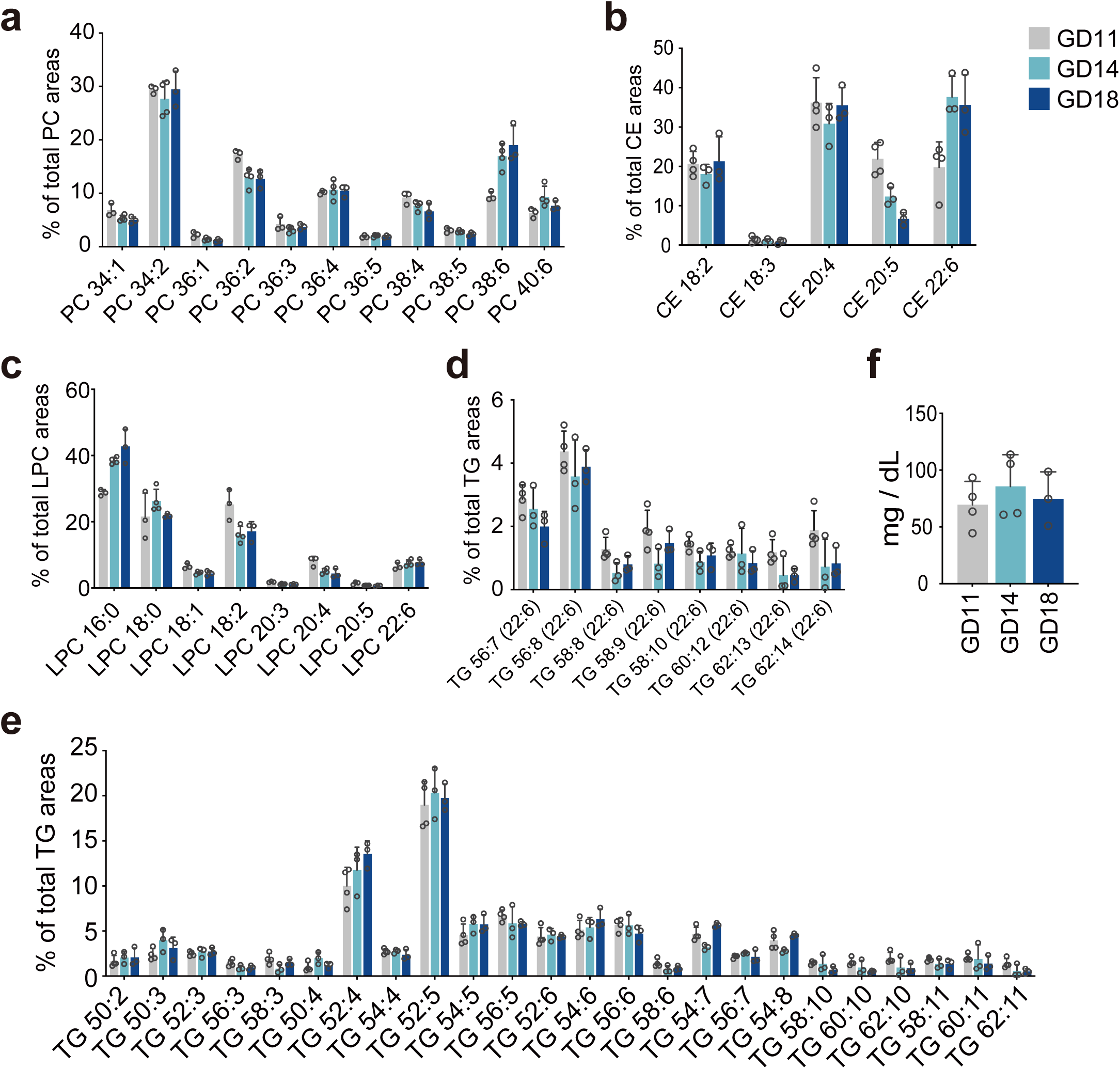
Serum lipid profiles of pregnant wild-type mice at gestational days 11 (GD11), GD14, and GD18. (**a**-**c**) Fatty acid compositions of phosphatidylcholine (PC) (**a**), cholesterol ester (CE) (**b**), and lysoPC (LPC) (**c**). The relative abundance in each lipid class is shown. PC species were classified by sum of carbon and double bond number of fatty acids at sn-1 and sn-2 positions. (**d**) The relative abundance of triglyceride (TG) species, classified by the sum of carbon and double bond number of three fatty acids, at GD11, GD14, and GD18. (**e**) The relative abundance of DHA-containing TG (DHA-TG), which contains at least one DHA in its structure. (**f**) Serum TG amounts of GD11, GD14, and GD18 mice. Each dot represents an individual mouse (**a** and **c**, n = 3, 4, and 3; **b**, **d**, and **e**, n = 4, 3, and 3; **f**, n = 4, 4, and 3 for GD11, 14, 18, respectively). Data are represented as means + SD.

**Supplementary Fig. 2.**
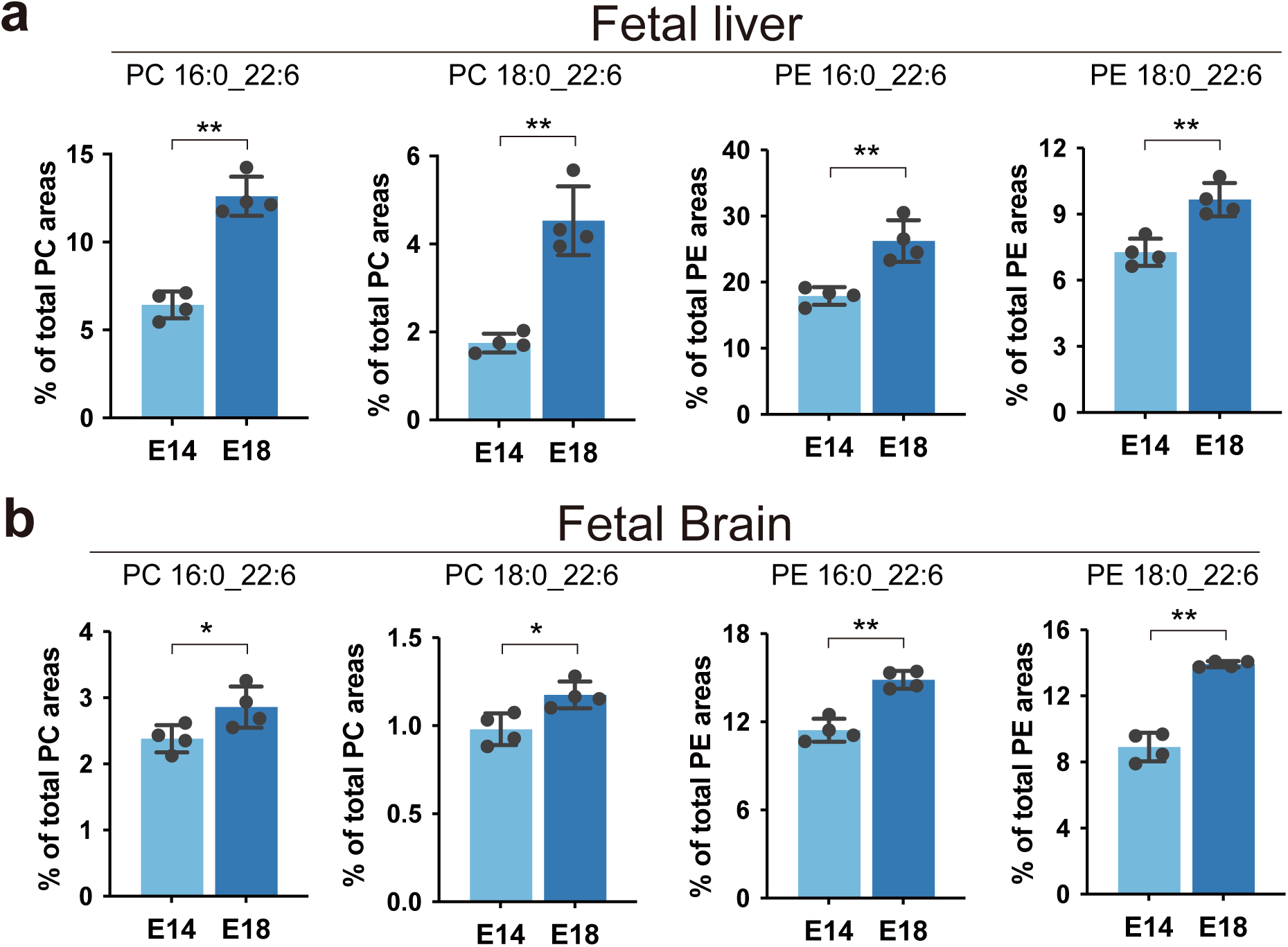
Levels of docosahexaenoic acid-phosphatidylcholine (DHA-PC) and DHA-phosphatidylethanolamine (DHA-PE) in the fetal liver and fetal brain of WT mice. (**a**-**b**) The relative amount of DHA-PC (PC 16:0_22:6 and PC 18:0_22:6) and DHA-PE (PE 16:0_22:6 and PE 18:0_22:6) in the fetal liver (**a**) and brain (**b**) at embryonic day 14 (E14) and E18. Each dot represents an individual mouse (n = 4 for each group). Data are presented as means ± SD. Statistical significance was determined using an unpaired t-test: \**P*<0.05 and \*\**P*<0.01.

**Supplementary Fig. 3.**
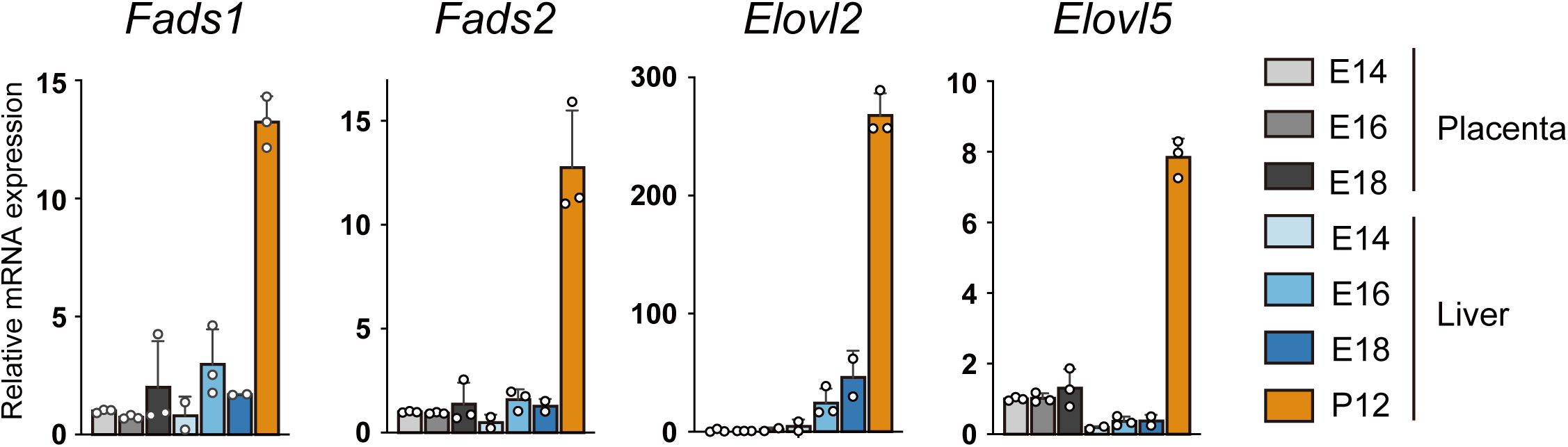
mRNA expression of enzymes involved in polyunsaturated fatty acid synthesis in the placenta and fetal liver. The relative expression of Fads1, Fads2, Elovl2, and Elovl5 mRNA in the placenta at embryonic day 14 (E14), E16, and E18 and the liver at E14, E16, E18, and postnatal day 12 (P12), normalized by Rplp0. Each dot represents an individual mouse (n = 3 for each group). Data are presented as means + SD.

**Supplementary Fig. 4.**
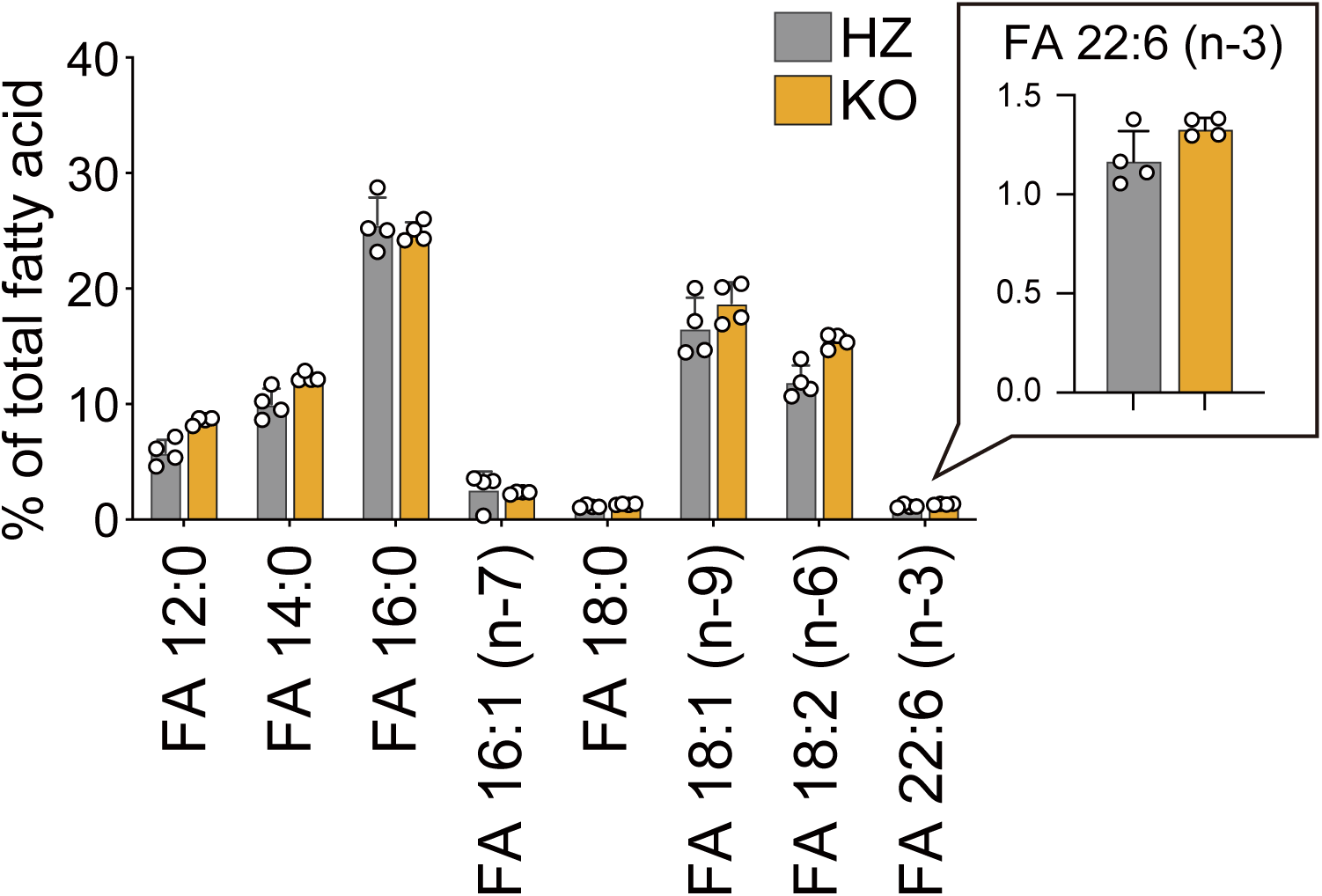
Fatty acid composition of breast milk. The relative amount of each fatty acid in the total lipid of the breast milk is shown. The relative amount of docosahexaenoic acid (DHA; represented as FA 22:6 (n-3)) is magnified in the boxed area. Breast milk was collected from the stomach of newborn (postnatal day 1) *Agpat3*-HZ and - KO mice. Each dot represents an individual mouse (n = 4 for each group). Data are represented as means + SD.

**Supplementary Fig. 5.**
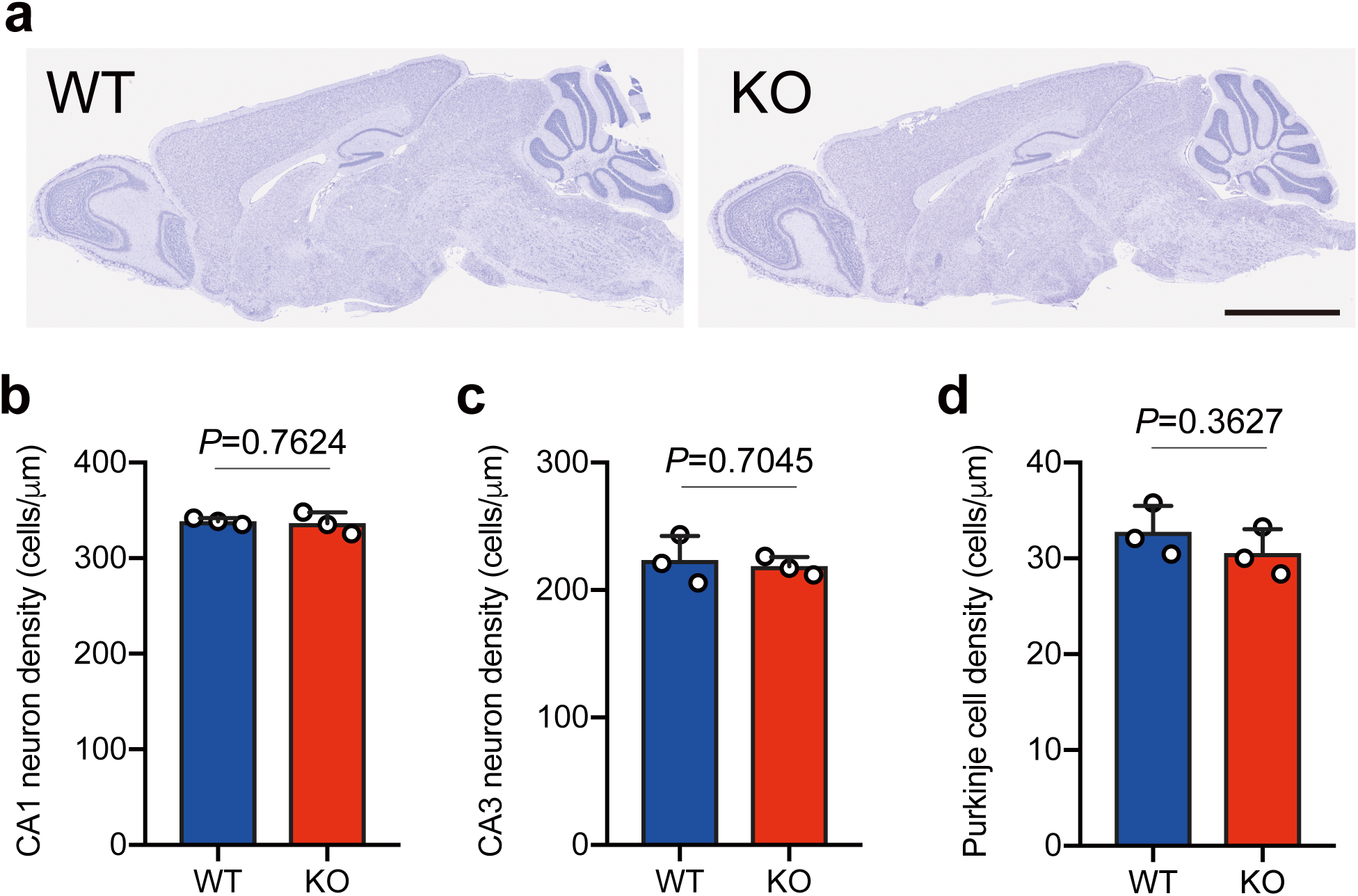
Histological analysis of *Agpat3*-KO mouse brain. (**a**) Representative Nissl staining images of 12-week-old male *Agpat3*-WT (left) and -KO (right) mouse brain. Scale bar: 2.5 mm. (**b**-**d**) Quantification of neuron density at hippocampal regions CA1 (**b**) and CA3 (**c**), and Purkinje cell density at cerebellum (**d**). Each dot represents an individual mouse (n = 3 for each group). Data are presented as means + SD. The *P*-values were determined using an unpaired t-test.

**Supplementary Fig. 6.**
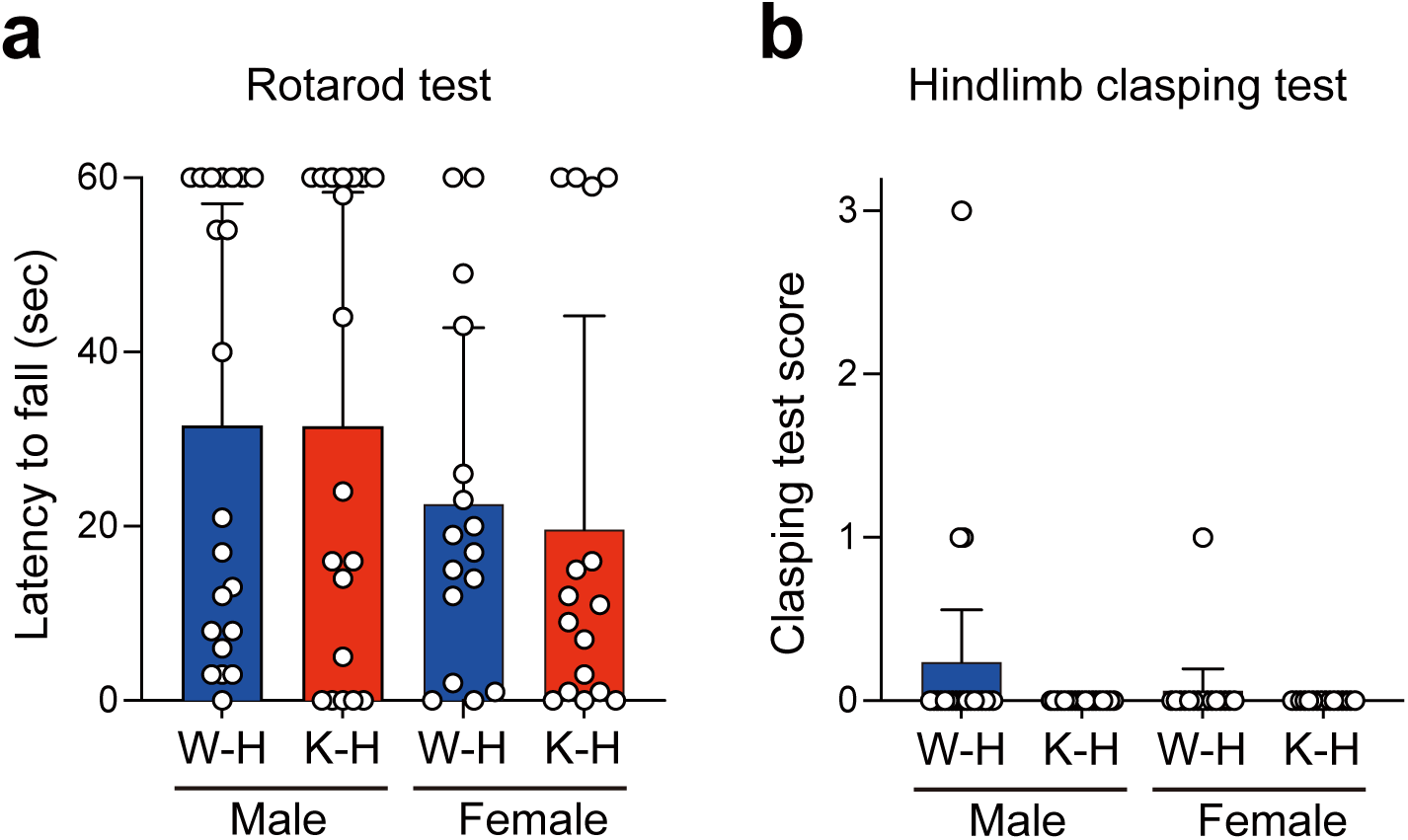
Behavioral phenotypes of *Agpat3*-HZ mice born from *Agpat3*-WT and -KO mice. (**a** and **b**) The results of rotarod (**a**), hindlimb clasping test (**b**). W-H and K-H indicate maternal–fetal *Agpat3* genotype (W, H, and K indicate WT, HZ, and KO, respectively). (**a**) The rotarod test was performed using 12-week-old mice. Falling latency of each mouse (n = 21, 19, 16, and 16 for male W-H, male K-H, female W-H, and female K-H, respectively) on a rotating rod at 10 rpm (60-s cutoff). (**b**) A hindlimb clasping test was performed using 4-week-old mice. Hindlimb clasping reflex was scored on a scale of 0–4 as described in the Methods section. Each dot in the bar graph showed the hindlimb clasping scores (n = 21, 19, 16, and 16 for male W-H, male K-H, female W-H, and female K-H, respectively). Data are presented as means ± SD. Statistical significance was determined using Bonferroni’ s multiple comparisons. Significant differences (*P* < 0.05) were not observed for all comparisons.

